# CRABS-ROC, A Respirometry Protocol For Overcoming Substrate Limitations, Reveals Excess Brain Mitochondrial Complex I Capacity

**DOI:** 10.1101/2025.07.23.666392

**Authors:** Naibo Zhang, Brian A. Roelofs, Evan A. Bordt, Boris Piskoun, Courtney L. Robertson, Brian M. Polster

## Abstract

Mitochondrial bioenergetic competency in cells is frequently assessed by the Mito Stress Test protocol, which includes uncoupler addition for evaluating respiratory capacity. The uncoupled oxygen consumption rate (OCR) is usually defined as maximal respiration, with little consideration of whether the measured rate is restricted by substrate supply. In this study, we show that the uncoupled OCR is substrate-limited in rat primary cortical neurons and isolated mouse forebrain synaptosomes. We use a different respirometry protocol we name CRABS-ROC (<u>C</u>omplex <u>R</u>espirometry <u>A</u>ssay <u>B</u>ypassing <u>S</u>ubstrate-<u>R</u>estricted <u>O</u>xygen <u>C</u>onsumption) that enables evaluation of individual electron transport chain (ETC) complex capacity using saturating levels of substrate to bypass this restriction. Applying CRABS-ROC to primary cortical neurons reveals >2-fold excess Complex I capacity beyond the uncoupled OCR of cells metabolizing glucose and pyruvate. Furthermore, we demonstrate that CRABS-ROC can expose a Complex I deficit in isolated harlequin mutant brain mitochondria that display wild-type levels of Complex I-substrate-linked respiration despite having about half the normal level of Complex I. Thus, CRABS-ROC should be broadly useful for studies on mitochondrial function because it can both reveal excess ETC capacity and unmask ETC alterations that may be missed by the most widely used methods.

## 1. Introduction

The energy storage molecule ATP is essential for brain function. Mitochondria couple electron transport to ATP generation through a series of membrane-embedded proton pumping complexes that produce an electrochemical gradient with potential energy [1]. Oxygen is the final electron acceptor in this oxidative phosphorylation process that uses the proton-motive force of the gradient to drive ADP to ATP phosphorylation. As cellular respiration consists primarily of mitochondrial oxygen consumption, measurement of oxygen consumption rate (OCR) is a convenient way to evaluate the bioenergetic competency of mitochondria within cells. However, in healthy cells OCR is dictated by proton gradient utilization by the ATP synthase, which in turn is driven by the ADP/ATP ratio or “energy demand” [1]. Therefore, to assess the bioenergetic capabilities of the mitochondrial network in cells, it is necessary to either increase energy demand or remove this restriction.

Chemical uncouplers like carbonyl cyanide 4-(trifluoromethoxy)phenylhydrazone (FCCP) and 2,4-dinitrophenol are widely used as part of the Mito Stress Test to evaluate mitochondrial capacity because they accomplish the latter goal and are not cell-type specific [2, 3]. Uncouplers dissociate electron transport from ADP phosphorylation, which increases OCR by removing ATP synthase control [1]. Thus, the OCR in uncoupler-treated cells is usually considered as maximal respiration, reflecting the mitochondrial “respiratory capacity,” while the difference between uncoupler-stimulated OCR and basal OCR is thought to indicate the “spare” or “reserve” respiratory capacity of the cell [2, 3].

A drawback of this method for assessing mitochondrial capability, however, is that the uncoupled respiration rate depends not only on the intrinsic capacity of the mitochondrial electron transport chain (ETC) components, which includes the ETC protein complexes, but also on the rate of electron entry from its substrates, NADH and FADH_2_. While FADH_2_ is covalently bound to several enzymes that converge to feed electrons to coenzyme Q, NADH, the immediate substrate of Complex I (NADH:ubiquinone oxidoreductase; EC 7.1.1.2), is provided primarily by dehydrogenase enzymes of the tricarboxylic acid (TCA) cycle [1]. Pyruvate, the glycolysis end-product that enters the TCA cycle as acetyl-CoA, is cell permeable and can be added with uncoupler to bypass the glycolysis rate restriction of OCR [2, 3]. Nevertheless, the activity level of one or more NADH-yielding dehydrogenase enzymes may still be a rate-limiting factor for measuring the true capacity of Complex I, the main site of ETC electron entry. Consistent with this possibility, evidence suggests that pyruvate dehydrogenase (PDH; EC 1.2.4.1), which is regulated by post-translational modifications that include phosphorylation and lipoylation, exerts the major control over mitochondrial respiration rate in many cell types [4].

To help overcome the limitation of evaluating mitochondrial bioenergetic competency using uncouplers, in this study we employ a simple respirometry protocol we name CRABS-ROC (<u>C</u>omplex <u>R</u>espirometry <u>A</u>ssay <u>B</u>ypassing <u>S</u>ubstrate-<u>R</u>estricted <u>O</u>xygen <u>C</u>onsumption) that enables direct interrogation of the capacity of individual complexes within the ETC in parallel to classical cellular respiratory capacity measurements that are influenced by substrate restrictions. We show that in addition to cells, the protocol is readily applied to isolated mitochondria or tissue homogenate, requiring only high nanogram to low microgram amounts of protein, respectively. By applying CRABS-ROC to harlequin (Hq) mutant mouse mitochondria that show ∼50% respiratory chain Complex I deficiency [5-8] due to insufficiency of the mitochondrial import protein apoptosis-inducing factor (AIF) [9-12], we demonstrate that substrate supply limits the respiration of brain mitochondria, which have excess Complex I capacity to support oxygen consumption.

## 2. Materials and Methods

### 2.1. Materials

Saponin (catalogue #S7900), alamethicin (catalogue #A4665), cytochrome *c* (catalogue #C2506), NADH (catalogue #10128023001) and pyruvate (catalogue #P2256) were from Sigma-Aldrich (St. Louis, MO). The bicinchoninic acid (BCA) assay was purchased from Thermo Fisher (Waltham, MA). Seahorse XF24 and XFe96 FluxPaks were acquired from Agilent Technologies (Santa Clara, CA) or Seahorse Bioscience (North Billerica, MA). Other reagents were obtained from Sigma-Aldrich unless noted otherwise. NADH and pyruvate were prepared fresh from powder immediately prior to experiments.

### 2.2. Animals

Male harlequin (Hq) mutant mice (strain name B6CBACa *A*^*w-J*^/*A*-*Aifm1*^*Hq*^/J) and age-matched male wild-type (WT) controls on the same background were acquired from the Jackson Laboratory (Bar Harbor, ME). Mice were kept on a 12 hour light/dark cycle, fed ad libitum, and housed according to standard animal care protocols. Embryonic Day 18 Sprague-Dawley rats from Charles River (Wilmington, MA) were used for the preparation of mixed-sex primary rat cortical neuron cultures. Postnatal day 18 male rats were used for the preparation of brain homogenates. All protocols were approved by the Institutional Animal Care and Use Committee (IACUC) and were in accordance with the *International Guiding Principles for Biomedical Research Involving Animals*, as issued by the Council for the International Organizations of Medical Sciences.

### 2.3. Preparation of primary neuron cultures

E18 rat cortices were dissociated using trypsin to prepare primary rat cortical neurons as previously described [13, 14]. Cells were plated in V7-PS or V28-PS microplates (Agilent Technologies) at a density of 90,000 or 100,000 cells/well (0.32 cm^2^) in Neurobasal medium (100 μl) containing B27 supplement (2%), fetal bovine serum (10%), L-glutaMAX (0.5 mM), streptomycin (100 μg/ml), and penicillin (100 IU/ml). Medium was replaced with 0.5 ml of the same medium but lacking fetal bovine serum at two hours post-plating. Cytosine arabinofuranoside (5 μM) was added on days *in vitro* (DIV) 4 to inhibit glial proliferation. Without removing any medium, fresh Neurobasal medium (0.2 ml) containing supplements was added to each well on DIV 6. Neurons were maintained at 37ºC in a humidified atmosphere of 95% air/5% CO_2_ and used for experiments at 6-15 DIV.

### 2.4. Preparation of brain homogenate

Following euthanasia, the brain was rapidly removed, and frontal and occipital brain tissue was dissected with a razor blade on an acrylic brain matrix previously cooled in ice. Using sharp scissors, the tissue was then rapidly minced in MS+EGTA buffer consisting of 225 mM mannitol, 75 mM sucrose, 1 mM ethylene glycol-bis(β-aminoethyl ether)-N,N,N′,N′-tetraacetic acid (EGTA), 1 mg/ml fatty acid-free bovine serum albumin (BSA), and 5 mM HEPES, pH 7.4. The minced tissue was then washed with MS+EGTA buffer and homogenized in the same buffer using a glass Dounce homogenizer with six strokes of pestle A (loose-fitting), followed by six strokes of pestle B (tight-fitting). The homogenized sample was centrifuged at 1000×*g* for 10 minutes at 4 ºC to remove nuclei and unbroken cells and then stored at -80 ºC until used for OCR measurements.

### 2.5. Isolation of mitochondria and synaptosomes

A Percoll™ density gradient was used to isolate non-synaptosomal brain mitochondria from 2-3 Hq male mice and from 2-3 male wild-type littermates in parallel as previously described [15]. Mouse forebrain synaptosomes were isolated from male C57BL6/J mice as described [7]. Mitochondrial and synaptosomal protein concentrations were determined by the Bio-Rad (Hercules, CA) Bradford method or by the BCA method (Thermo Fisher). Mitochondria and synaptosomes were stored on ice and used for OCR measurements on the day of isolation and then frozen at -80º C. Following thawing on a later date, the same mitochondria were used for OCR measurements by the CRABS-ROC protocol.

### 2.6. Seahorse respirometry – neurons and synaptosomes

Neuronal [16] and synaptosomal [17] OCR measurements were made using a Seahorse XF24 Extracellular Flux Analyzer (Agilent Technologies) as previously described. Artificial cerebrospinal fluid (aCSF) assay medium consisted of 120 mM NaCl, 3.5 mM KCl, 1.3 mM CaCl_2_, 0.4 mM KH_2_PO_4_, 1 mM MgCl_2_, 4 mg/ml fatty acid-free BSA, and 5 mM HEPES, pH 7.4. Glucose was additionally present at 15 mM unless indicated otherwise. Cells or synaptosomes were incubated in aCSF assay medium for one hour prior to measurements in a CO_2_-free incubator at 37°C. For some experiments, cells were incubated in aCSF for 45 minutes prior to this step in a 95% air/5% CO_2_ incubator at 37°C, followed by an additional media change to fresh aCSF.

For the experiments in which the permeabilizing agents saponin and alamethicin were employed, the aCSF contained an increased concentration of HEPES (20 mM) because greater buffer capacity was required. Also, for those experiments the neurons were plated on V28-PS plates, which have a higher measurement volume (28 μl) than that of V7-PS plates (7 μl). The greater volume prevented excessive oxygen depletion during the measurement period due to the high OCR of NADH-stimulated respiration.

### 2.7. Seahorse respirometry – homogenate or isolated mitochondria

Brain homogenate and isolated brain mitochondria were diluted in MAS assay medium to concentrations of 0.4 mg/ml and 0.02 mg/ml, respectively. The MAS medium consisted of 220 mM mannitol, 70 mM sucrose, 5 mM KH_2_PO_4_, 5 mM MgCl_2_, 1 mM EGTA, and 2 mM HEPES, pH 7.4. Twenty μl of the diluted suspensions were added to XFe96 plates (Agilent Technologies) for final amounts of 8 μg brain homogenate and 0.4 μg brain mitochondria, respectively, while four wells on each plate received 20 μl of MAS medium to be used as a temperature control. The XFe96 plates were centrifuged at 2,000×*g* for 5 minutes at 4 ºC without braking to adhere mitochondria to the bottom of the wells, and then 160 μl of MAS assay medium was added to each well. OCR was measured using a Seahorse XFe96 Extracellular Flux Analyzer (Agilent Technologies) as previously described [18].

### 2.8. Statistics

OCR were compared by paired *t* test for two groups and by two-way ANOVA with repeated measures for more than two groups using GraphPad Prism version 10.4.1 (GraphPad, Boston, MA). Following the ANOVA, the Šídák post hoc test was used for pairwise comparisons. *P*<0.05 was considered significant.

## 3. Results

### 3.1. The uncoupled oxygen consumption rate in the Mito Stress Test is limited by substrate supply

Mitochondrial respiratory capacity is usually assessed following uncoupler addition as a component of the widely used Mito Stress Test protocol [2, 19]. The protocol also includes the prior addition of the ATP synthase inhibitor oligomycin to estimate mitochondrial coupling. We applied the Mito Stress Test protocol to primary rat cortical neurons metabolizing a physiological concentration of glucose (3 mM), which supplies the glycolysis end product pyruvate to mitochondria, and then added an excess amount of pyruvate (10 mM) [20] to determine whether maximal respiration had been obtained upon addition of the uncoupler FCCP. Significantly greater OCR was obtained after pyruvate addition compared to the uncoupled OCR of neurons metabolizing only physiological glucose (Fig. 1A, B). Most bioenergetics studies are done in the presence of supraphysiological glucose concentrations. To determine if the substrate limitation is overcome at a typical supraphysiological glucose concentration, we repeated the experiment using neurons incubated with 15 mM glucose and obtained the same results (Fig. 1A, B). Together, these results suggest that mitochondrial substrate supply is rate-limiting in neurons.

**Fig. 1.**
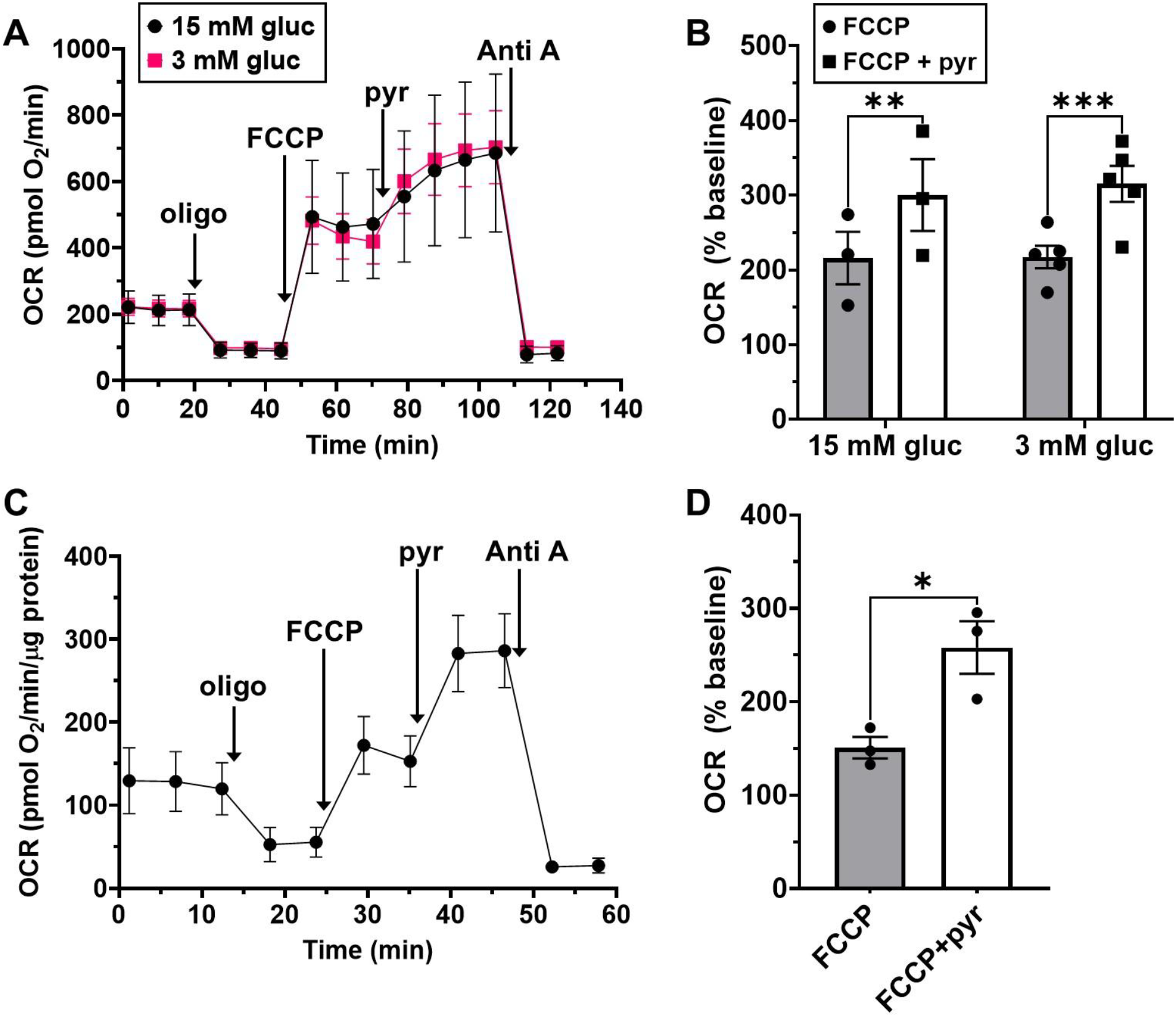
Uncoupled respiration is limited by substrate availability. **A**. The oxygen consumption rate (OCR) of primary cortical neurons at baseline and following the sequential addition of oligomycin (oligo, 0.5 μg/ml), FCCP (3 μM), pyruvate (pyr, 10 mM), and antimycin A (Anti A, 1 μM). The results are mean ± SEM from biological replicates. The glucose (gluc) concentration was 3 mM (*n*=3) or 15 mM (*n*=5), as indicated. **B**. The maximal OCR following the FCCP and pyr additions (mean ± SEM). \**p*<0.01; \*\*\**p*<0.001. **C**. The OCR of forebrain synaptosomes at baseline and following the sequential addition of oligo (2 μg/ml), FCCP (4 μM), pyr (10 mM), and Anti A (10 μM). The results are mean ± SEM from *n*=3 biological replicates. **D**. The maximal OCR following the FCCP and pyr additions (mean ± SEM). \**p*<0.05

The mitochondrial reliance of cells cultured in glucose is thought to be low, whereas synaptic function is thought to be highly dependent on mitochondrial ATP supply. Therefore, we acutely isolated pre-synaptic terminals (“synaptosomes”) from mouse forebrain to further investigate substrate limitations. Pyruvate addition led to a significant stimulation of the uncoupled OCR of synaptosomes metabolizing glucose, very similar to the enhancement observed in cells (Fig. 1C, D). Previously, we showed that pyruvate also enhances the uncoupled respiration measured in mouse organotypic hippocampal slices, which consist of multiple brain cell types in their native architecture [21]. Therefore, the results from a variety of neural models suggest that substrate supply is rate-limiting when measuring mitochondrial respiratory capacity using uncoupler in the Mito Stress Test.

### 3.2. Complex I has excess capacity that is obscured in neurons metabolizing glucose

NADH, not pyruvate, is the direct Complex I substrate. Therefore, it is possible that the provision of NADH was still rate-limiting in measuring respiratory capacity with the uncoupler/pyruvate combination. However, NADH is not cell permeable and, thus, cannot be readily incorporated into the Mito Stress Test protocol. Previously, we optimized Seahorse cell-based respirometry for measuring respiration by permeabilized cells [16]. The assay uses saponin to deliver a defined ETC substrate combination to mitochondria by selectively permeabilizing the plasma membrane. Subsequently, we showed that Complex I-III-IV linked activity could be measured using Seahorse respirometry by additionally adding the bacterial pore forming peptide alamethicin to transport the Complex I substrate NADH across the mitochondrial inner membrane [22]. This assay required the provision of exogenous cytochrome *c* to replace the endogenous cytochrome *c* released by alamethicin.

To evaluate whether primary rat neurons have excess Complex I capacity, we employed the permeabilized cell assay and a Complex I-III-IV linked activity assay in parallel while also comparing OCR to intact neurons treated with FCCP and an overabundance of the cell permeable Complex I substrate pyruvate (Fig. 2). The three different compound injection protocols were done simultaneously on a single plate. Whereas the ADP-stimulated respiration of permeabilized cells provided with the Complex I-linked substrates pyruvate and malate was similar to that measured in intact cells treated with FCCP plus pyruvate (Fig. 2), a >2-fold higher OCR was measured when alamethicin was used to deliver the immediate Complex I substrate NADH (Fig. 2). These results revealed that neuronal Complex I capacity is obscured when tricarboxylic acid (TCA) cycle dehydrogenase substrates drive respiration.

**Fig. 2.**
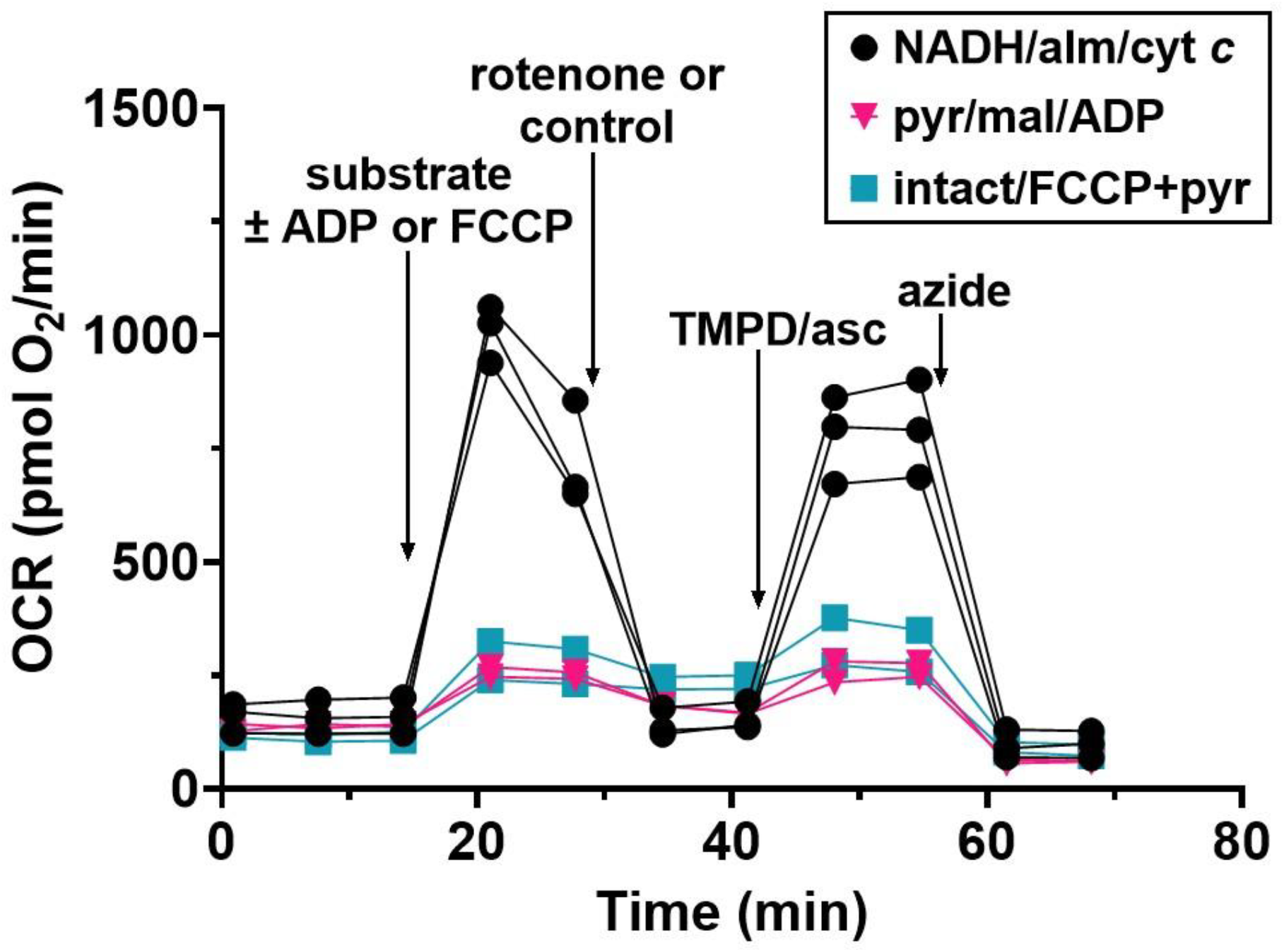
NADH delivery to mitochondria with primary rat cortical neurons reveals excess Complex I capacity. Three baseline O_2_ consumption rate (OCR) measurements were made and then one of the following three cocktails was injected as indicated: 0.5 mM NADH + 40 μg/ml alamethicin + 100 μM cytochrome *c* (“NADH/alm/cyt *c*”), 5 mM pyruvate + 5 mM malate + 1 mM ADP (“pyr/mal/ADP”), or 3 μM FCCP + 10 mM pyruvate (“intact/FCCP+pyr”). Saponin (sap, 3 μg/ml) plus EGTA (5 mM) and K_2_PHO_4_ (3.6 mM) were additionally present in the “NADH/alm/cyt c” and “pyr/mal/ADP” injections. Rotenone (1 μM) or control, TMPD (0.1 mM) + ascorbate (asc, 10 mM), and sodium azide (5 mM) were then sequentially injected as indicated. Only the NADH/alm/cyt *c* group received rotenone; the other groups received an aCSF control injection. The results are technical replicates of one experiment and are representative of two (pyr/mal/ADP group) to three (other groups) biological replicates.

### 3.3. CRABS-ROC unmasks a Complex I activity deficit in harlequin mutant mitochondria

Next, we sought to apply the Complex I-III-IV-linked activity assay to a model of genetic Complex I deficiency, the harlequin mouse, while also incorporating measurement of Complex II activity. We named this modified assay protocol “CRABS-ROC” to reflect its ability to bypass endogenous substrate-restricted oxygen consumption by providing the respective Complex I and II substrates NADH and succinate in excess. Harlequin mice harboring a mutation in the gene encoding AIF [23], a component of the mitochondrial disulfide relay protein import pathway [9-12], have about a 2-fold reduction in assembled Complex I [5-8]. Unexpectedly, we [7] and others [6] found that despite this Complex I deficiency, isolated harlequin mouse brain mitochondria show a wild-type ADP-stimulated respiration level when oxidizing the Complex I-linked substrates glutamate and malate. This finding suggests that even in isolated mitochondria, the full Complex I capacity may be obscured when Complex I-linked TCA cycle dehydrogenase substrates are used.

CRABS-ROC can be applied to frozen tissue samples [18], giving us the opportunity to re-evaluate Complex I activity in the stored isolated brain mitochondria samples from our prior study. As the cytochrome *c* and alamethicin concentrations in the Respirometry In Frozen Samples (RIFS) protocol [18] differ from those we optimized for neurons in cell culture under different experimental conditions [22], we first re-titrated cytochrome *c* and alamethicin using RIFS on freeze-thawed rat brain homogenate. We found that concentrations of 100 μM cytochrome *c* (Fig. 3A) and 40 μg/ml alamethicin (Fig. 3B) yielded maximal respiration whereas for brain homogenate, the previously employed concentrations of 8 μM (10 μg/ml) cytochrome *c* (Fig. 3A) and 10 μg/ml alamethicin (not shown) [18] were suboptimal.

**Fig. 3.**
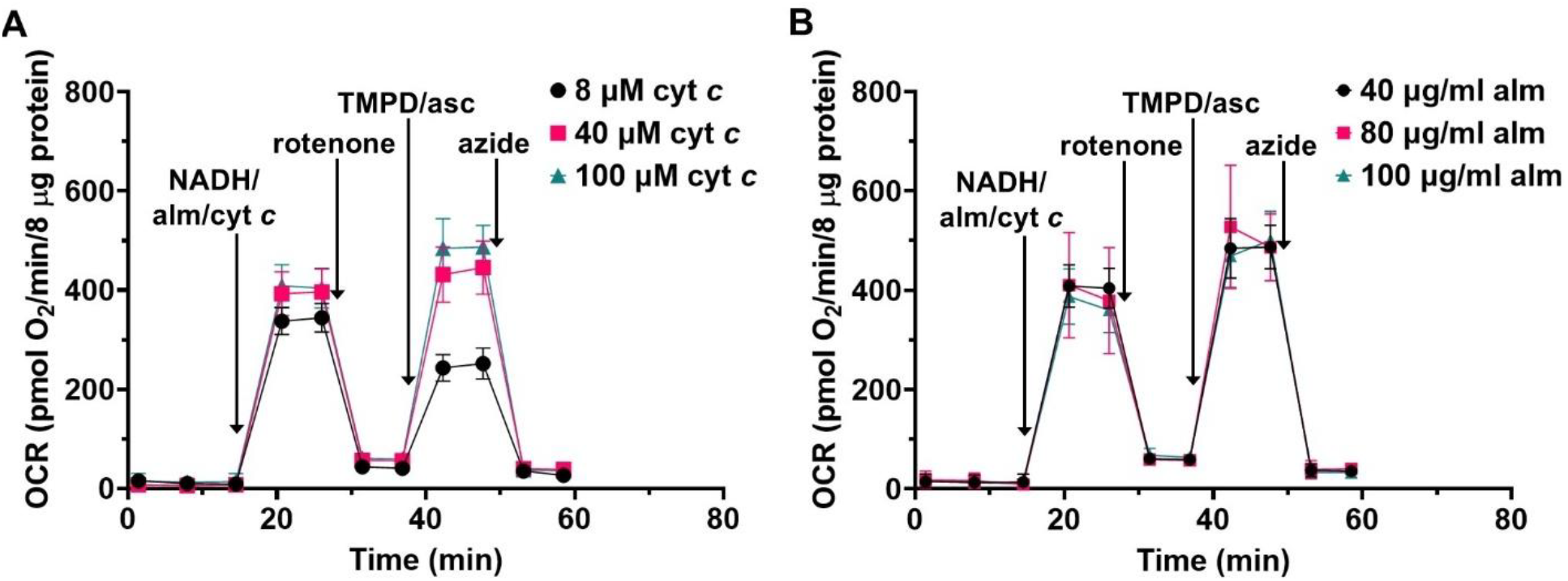
Titrations of cytochrome *c* (cyt *c*) and alamethicin (alm) for the Complex Respirometry Assay Bypassing Substrate-Restricted Oxygen Consumption (CRABS-ROC) protocol. **A**. Oxygen consumption rate (OCR) measurements from brain homogenate. NADH (1 mM) was added with the pore-forming peptide alm (40 μg/ml) and cyt *c* to initiate Complex I dependent respiration, followed by the Complex I inhibitor rotenone (2 μM), the Complex IV substrate TMPD (0.4 mM) with ascorbate (asc, 2 mM), and the Complex IV inhibitor sodium azide (50 mM). The cyt *c* concentration was varied as shown. The results are mean ± SD, *n*=6-8 technical replicates per group. **B**. OCR measurements from brain homogenate. The same additions were made as in A, but the cyt *c* concentration was 100 μM and the alm concentration was varied as shown. The results are mean ± SD, *n*=8 technical replicates per group.

Next, WT and Hq mitochondria, which displayed equivalent ADP-stimulated respiration when oxidizing the Complex I-linked substrates malate and glutamate (Fig. 4A), were subjected to an optimized CRABS-ROC protocol consisting of the sequential additions of NADH, rotenone, succinate, and combined TMPD/ascorbate. By including some wells in which sodium azide was injected together with TMPD/ascorbate, this protocol allows for the measurement of Complex I, Complex II, and Complex IV activities in individual samples (Fig. 4B). As expected, we found no difference in Complex IV-dependent respiration in Hq mitochondria compared to WT. However, the Complex I-dependent OCR measured upon NADH addition was significantly lower in Hq mitochondria compared to WT (Fig. 4C). The CRABS-ROC protocol also revealed a strong trend (*p*=0.06) toward increased Complex II-dependent OCR in Hq mitochondria (Fig. 4D) and the ratio of Complex I to Complex II-dependent OCR was decreased by over two-fold (Figure 4E). Thus, CRABS-ROC unmasked a Complex I activity deficit that multiple studies [6, 7] failed to reveal by conventional respirometry using Complex I-linked substrates. It also revealed a likely adaptive increase in Complex II activity.

**Fig. 4.**
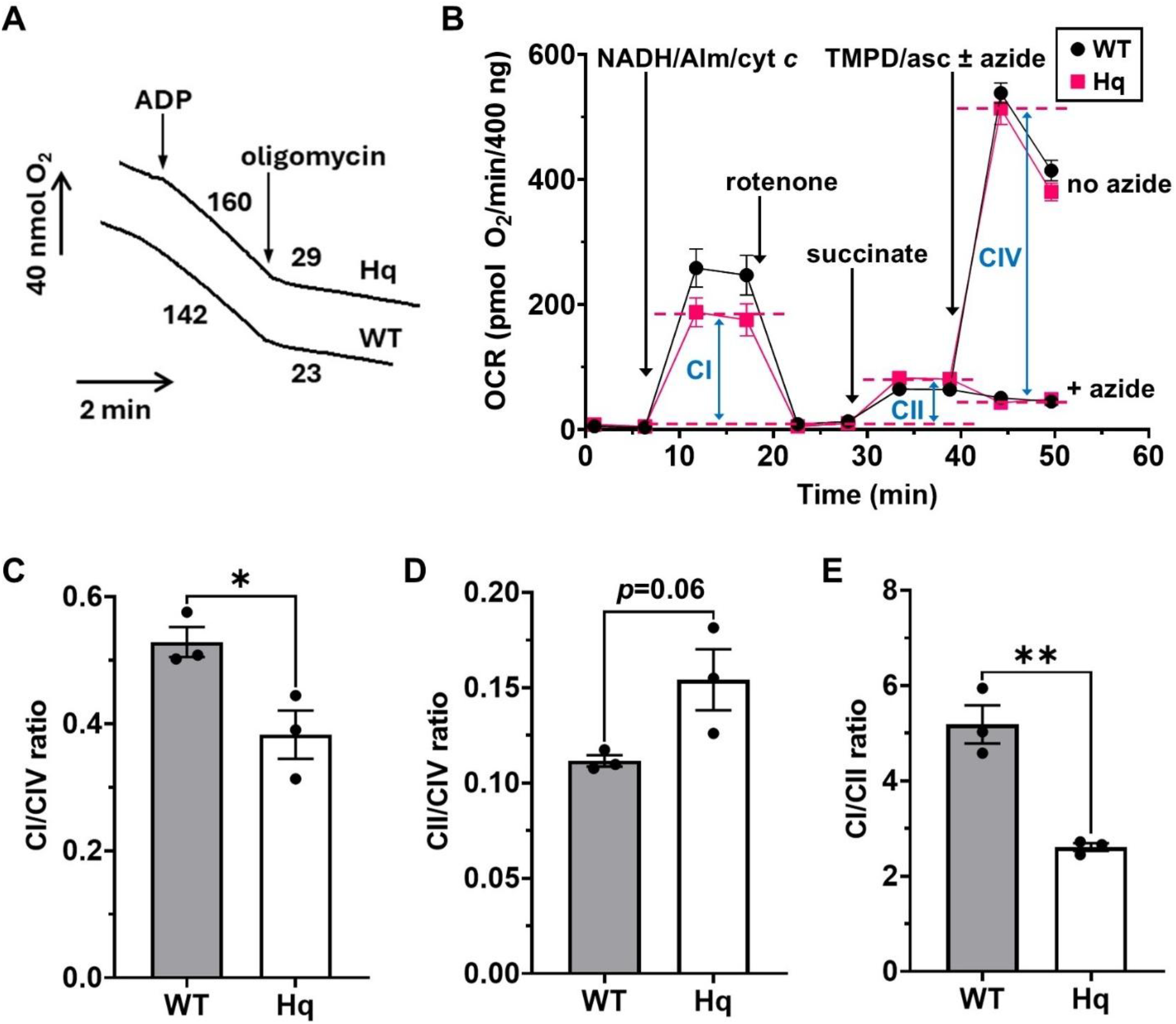
CRABS-ROC unmasks Complex I deficiency in AIF-depleted harlequin (Hq) brain mitochondria. **A**. Reprinted with permission from Fig. 1C of [7]. ADP (1 mM)-stimulated oxygen consumption was measured using a Clark-type oxygen electrode at 30°C for wild-type (WT) and Hq brain mitochondria (0.25 mg/ml) incubated in a potassium chloride-based assay medium. Oligomycin (3 μg/ml) was then added. Oxygen consumption rates (OCR) are shown in nmol O_2_/min/mg mitochondrial protein; additional experimental details can be found in [7]. **B**. OCR measurements from WT and Hq brain mitochondria. NADH (1 mM) was added with the pore-forming peptide alamethicin (alm, 40 μg/ml) and cytochrome *c* (cyt c, 100 μM), followed by rotenone (2 μM), succinate (5 mM), and TMPD (0.4 mM) plus ascorbate (asc, 2 mM). Sodium azide (50 mM) was added with TMPD/asc when indicated. The OCR subtractions used to calculate Complex I-dependent (I), Complex II-dependent (II), and Complex IV-dependent (IV) OCR are shown graphically for the Hq trace and were done in the same way for the WT trace, with calculations using the first data point following each injection. The results are mean ± SEM from *n*=3 biological replicates. **C**. The ratio of Complex I-dependent to Complex IV-dependent OCR. **D**. The ratio of Complex II-dependent to Complex IV-dependent OCR. **E**. The ratio of Complex I-dependent to Complex II-dependent OCR. The results in C-E are mean ± SEM, *n*=3. \**p*<0.05; \*\**p*<0.01

## 4. Discussion

Over the past two decades, mitochondrial bioenergetics research has expanded rapidly, largely driven by the introduction of the Seahorse Extracellular Flux Analyzer. This technology popularized the Mito Stress Test, a respirometry injection protocol that became the standard protocol in bioenergetics studies [2, 19]. However, in many—if not most studies—this was the only protocol employed, and potential limitations related to substrate availability were often overlooked.

Our experiments employing exogenous pyruvate reveal that the Mito Stress Test significantly underestimates mitochondrial respiratory capacity in neurons due to insufficient substrate provision. This limitation may also extend to other cell types. Using a complementary respirometry protocol we named CRABS-ROC, we demonstrated that substrate availability not only underestimates respiratory capacity, but it also masks the capacity of Complex I in both intact cells and isolated mitochondria. This suggests that maximal respiration is constrained by the rate at which tricarboxylic acid (TCA) cycle dehydrogenases supply NADH to the electron transport chain (ETC)—a process that is tightly regulated by enzymes such as PDH [4].

Importantly, the CRABS-ROC protocol was able to uncover a Complex I activity deficit in harlequin mutant mitochondria that went undetected by a conventional respirometry protocol, despite an approximately 50% reduction in most Complex I proteins [6, 7]. Previous studies required titration with Complex I inhibitors to functionally demonstrate the deficiency in harlequin mitochondria [6, 7]. This highlights the enhanced sensitivity of CRABS-ROC in detecting subtle bioenergetic impairments that may be present in many disease models. The existence of excess Complex I capacity likely explains why both harlequin mutant mice and *Ndufa1*^S55A^ mutant mice—each exhibiting ∼50% Complex I deficiency—maintain a relatively normal lifespan [23, 24], with female *Ndufa1* mice even showing a modest lifespan extension relative to WT controls [24].

Our observation of excess Complex I capacity has important implications for disease biology. Loss of Complex I function is implicated in the pathogenesis of Parkinson’s disease [25-27], a late-onset neurodegenerative disorder primarily affecting the elderly [28]. Exposure to environmental toxins that inhibit Complex I are linked to the sporadic form of Parkinson’s disease [29-31]. Our finding that neurons possess substantial excess Complex I capacity may help explain in part the disease’s typically late onset and slow progression, as a threshold of cumulative toxin exposure may be required prior to the manifestation of clinical symptoms.

While the CRABS-ROC protocol offers enhanced sensitivity for detecting mitochondrial respiratory deficits, it has several limitations. To facilitate Complex I substrate delivery, the protocol compromises the integrity of the inner mitochondrial membrane. As a result, it is less suitable for studies investigating mitochondrial structural dynamics, such as those involving alterations in fusion, fission, or membrane lipid composition. Additionally, CRABS-ROC relies on the addition of exogenous substrates and cytochrome c—an intermembrane space ETC component—at concentrations that may exceed physiological levels. Consequently, the protocol is better suited for assessing maximal ETC enzyme capacities rather than physiological ETC activity. Therefore, CRABS-ROC should not be viewed as a replacement for the Mito Stress Test, but rather as a complementary approach that can be run in parallel with assays involving intact or mitochondrial-outer-membrane-permeabilized cells.

## 5. Conclusions

Our study highlights the limitations of the widely used Mito Stress Test protocol in accurately assessing mitochondrial bioenergetic capacity due to substrate constraints. To address this, we introduced the CRABS-ROC protocol—a complementary approach that assesses the capacity of individual respiratory chain components and enables detection of hidden ETC deficits. Our findings emphasize the importance of considering substrate availability in bioenergetics research, especially when investigating mitochondrial dysfunction in disease models where the impairments may be subtle. This approach could also have translational applications in understanding functional aspects of human Complex I deficiencies.

## Abbreviations

AIF: apoptosis-inducing factor
alm: alamethicin
Anti A: antimycin A
asc: ascorbate
BCA: bicinchoninic acid
BSA: bovine serum albumin
CRABS-ROC: Complex Respirometry Assay Bypassing Substrate-Restricted Oxygen Consumption
cyt *c*: cytochrome *c*
DIV: days *in vitro*
EGTA: ethylene glycol-bis(β-aminoethyl ether)-N,N,N′,N′-tetraacetic acid
ETC: electron transport chain
FCCP: carbonyl cyanide 4-(trifluoromethoxy)phenylhydrazone
Hq: harlequin
mal: malate
OCR: oxygen consumption rate
PDH: pyruvate dehydrogenase
pyr: pyruvate
RIFS: Respirometry In Frozen Samples
TCA: tricarboxylic acid
WT: wild-type

## Acknowledgements

The authors thank Dr. Nagendra Yadava for helpful discussions and expert advice.

## CRediT authorship contribution statement

**Naibo Zhang:** Conceptualization, Data curation, Formal analysis, Investigation, Methodology, Roles/Writing - original draft, Writing - review & editing. **Brian A. Roelofs:** Conceptualization, Data curation, Formal analysis, Investigation, Methodology, Writing - review & editing. **Evan A. Bordt:** Conceptualization, Data curation, Formal analysis, Investigation, Methodology, Writing - review & editing. **Boris Piskoun:** Investigation, Writing - review & editing. **Courtney Robertson:** Conceptualization, Resources, Writing - review & editing. **Brian M. Polster:** Conceptualization, Data curation, Formal analysis, Funding acquisition, Investigation, Methodology, Project administration, Resources, Supervision, Roles/Writing - original draft, Writing - review & editing.

## Availability of data and materials

Not applicable.

## Declaration of Generative AI and AI-assisted technologies in the writing process

During the preparation of this work the authors used Microsoft Copilot to improve the wording of the Discussion and Conclusion sections. After using this tool, the authors reviewed and edited the content and take full responsibility for the content of the publication.

